# Predicting PROTAC-targeted Degradation and Designing Androgen Receptor Degraders with AiPROTAC

**DOI:** 10.1101/2025.02.26.640266

**Authors:** Li Zhang, Renhong Sun, Xia Li, Chaowei Ren, Yahui Long, Wenjuan Tan, Xiaolong Wang, Ke Zou, Xiaobao Yang, Min Wu, Xiaoli Li, Xing Chen

## Abstract

Proteolysis-targeting chimeras (PROTACs), a pioneering class of heterobifunctional ligands, have emerged as transformative tools in combating cancer and immune-related diseases due to their ability to target previously undruggable proteins and overcome drug resistance. Accurate assessment of the degradation potential of PROTACs is essential for advancing their therapeutic applications. However, the development of robust predictive models is hindered by the scarcity of large-scale datasets and domain-specific tools, as well as the underutilization of existing unlabeled data. To address these challenges, we developed an innovative method called AiPROTAC to predict the degradation capacity of PROTACs. In particular, AiPROTAC integrates graph augmentation, message-passing enhanced encoders and cross-attention mechanisms within a contrastive learning framework. Moreover, AiPROTAC leverages a curated specialized dataset combined with the PROTAC-DB dataset to enhance prediction accuracy. Experimental results across two datasets and six evaluation metrics demonstrate that AiPROTAC consistently outperforms state-of-the-art models. Further case studies underline its superior sensitivity and reliability. Notably, AiPROTAC facilitated the design of a novel androgen receptor (AR) degrader, PROTAC GT19, which achieved enhanced in vitro degradation compared to Bavdegalutamide (ARV-110). This advancement highlights our AiPROTAC’s potential to accelerate PROTAC-based therapeutic development and optimization.

## 1 Introduction

As a groundbreaking drug discovery strategy, proteolysis-targeting chimeras (PROTACs), which induce selective protein degradation via the ubiquitin-proteasome system (UPS), have garnered significant attention since their inception in 2001^1^. Biochemically, PROTACs are heterobifunctional molecules consisting of two ligands connected by a chemical linker: one, the warhead, targets the protein of interest (POI), while the other, the E3 ligand, recruits an E3 ligase^2–4^. Unlike conventional inhibitors, which demand strong binding to the POI, PROTACs only require transient interactions to trigger POI ubiquitination and degradation^5, 6^. In this context, Fig. 1a illustrates the mechanism by which PROTACs hijack the UPS to induce POI degradation. Using adenosine triphosphate (ATP) to generate activated ubiquitin-adenylate, the E1 ubiquitin-activating enzyme first activates ubiquitin, which is then transferred to the catalytic cysteine of the E2 ubiquitin-conjugating enzyme via the transthioesterification reaction^7, 8^. While E3 ligase cannot typically ubiquitinate a distant POI^9^, the PROTAC brings the POI and E3 ligase into close proximity, forming a ternary complex and driving the transfer of ubiquitin from E2 to the exposed lysine on the POI^10^. This process is repeated, generating a poly-ubiquitin chain, which is recognized by the 26S proteasome and leads to the degradation of the POI into small fragments or amino acids^11, 12^.

**Fig. 1.**
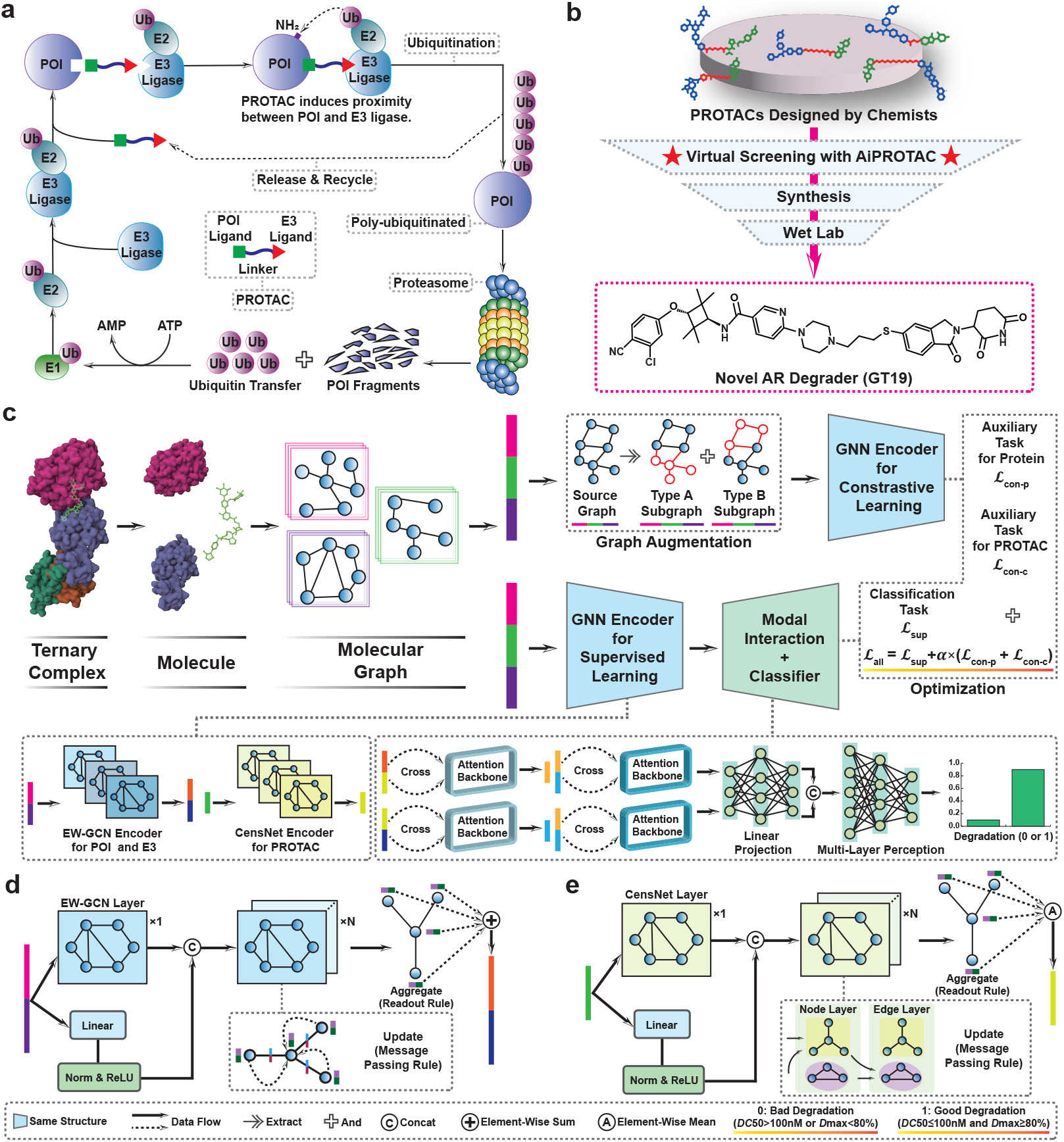
An overview of our AiPROTAC approach. **(a)**. The mechanism of PROTAC hijacking the UPS to induce POI degradation. In response to the PROTAC induction, POI and E3 ligase come into proximity, thereby initiating the ubiquitin transfer process, and polyubiquitination ultimately takes place on the POI. The ubiquitinated POI is recognized by proteasome and degraded into protein fragments. **(b)**. AI-based drug development paradigm in actual applications. An AI tool first screens PROTACs designed by chemists. Those that show strong degradation are then made and tested in the lab. **(c)**. The network architecture of AiPROTAC. Molecular graphs of POI, E3 ligase and PROTAC are simultaneously input into a classification network and two auxiliary networks. The classification network follows a standard encoder-decoder architecture, including GNN encoders, a modal interaction component and a classifier. Both auxiliary networks incorporate graph augmentation and GNN encoders. Note: The auxiliary tasks are only performed during training. **(d-e)**. The structure of EW-GCN encoder and CensNet encoder. Both are novel graph encoders enhanced with edge-based message passing.

Compared to traditional small molecule inhibitors that rely on occupancy-driven pharmacology, PROTACs utilize an event-driven mechanism of action with bifunctional small molecules, offering several distinct advantages: (i) PROTACs can modulate targets without classical hydrophobic binding pockets or those bound by endogenous molecules^13–15^. They also address proteins involved in protein-protein interactions, which are challenging for inhibitors that block catalytic activities or disrupt interfaces^16, 17^. (ii) PROTACs engage in a catalytic process, being released from the ternary complex after ubiquitination is completed^18^. This allows them to operate at low concentrations, reducing the risk of off-target effects. (iii) Unlike inhibitors that stabilize their target proteins and cause transcriptional upregulation^19, 20^, PROTACs drive the degradation of entire proteins via the proteasome, preventing accumulation and mitigating drug resistance from mutations around the binding pocket^21–24^. (iv) By exploiting sequence and conformational differences outside the catalytic core, PROTACs selectively degrade homologous proteins even with conserved active sites^25, 26^.

Despite over 20 years since the first proof-of-concept PROTAC, and the clinical progress of candidates like Bavdegalutamide (ARV-110) and Vepdegestrant (ARV-471), no PROTAC has yet been approved for market use^27^. This delay stems from the stringent requirements for druggable PROTACs, including effective degradation, specificity, cell permeability, pharmacokinetics, safety, synthetic feasibility, stability, and delivery^28–30^. These obstacles hinder the development of viable PROTAC drugs, with degradation capacity being the most critical factor in determining a PROTAC’s potential for further optimization and clinical application^31^. Traditional methods, which necessitate molecule synthesis followed by costly and time-consuming wet-lab experiments, contribute to high costs, long timelines, and low success rates^32^. Thus, alternative, more efficient approaches are urgently needed to streamline the process.

Building on recent advancements in artificial intelligence (AI) technologies^33, 34^ and high-quality PROTAC datasets^35^, several computational models have emerged to aid PROTAC drug discovery. Tools like DeLinker^36^, PROTAC-RL^37^, 3DLinker^38^, DiffLinker^39^, LinkerNet^40^, and DeepPROTACs^41^ exemplify this progress. For instance, DeLinker^36^ generates linkers between two given molecular fragments by integrating three-dimensional structural information into a graph-based encoder-decoder network. Zheng *et al*.^37^ combined an augmented transformer architecture with memory-assisted reinforcement learning in PROTAC-RL, which generates chemically feasible PROTACs with desired properties. 3DLinker, DiffLinker, and LinkerNet are atom-level PROTAC design models. 3DLinker^38^ uses a variational autoencoder to predict anchor atoms, and is capable of simultaneously generating linker graphs and structures. Both DiffLinker and LinkerNet employ a diffusion generation strategy that satisfies geometric equivariance. DiffLinker^39^ uniquely determines the number of atoms in the linker and can connect multiple molecular fragments. LinkerNet^40^ addresses the need to predefine the relative positions of two molecular fragments during linker generation. However, these models largely focus on linker design or PROTAC generation, with limited emphasis on predicting degradation efficacy. To overcome this, Li *et al*.^41^ proposed DeepPROTACs, a model that employs separate neural network modules to process distinct components of ternary complexes. While it shows promising performance, its predictive accuracy still has room for improvement, as it did not leverage the unlabeled data for model training. Moreover, its application in drug development pipelines remains largely unexplored.

To address the above challenges, we developed AiPROTAC, a novel deep graph model tailored to predict the degradation capacity of PROTACs. AiPROTAC integrates graph augmentation, two message-passing enhanced encoders and cross-attention mechanisms within a unified framework of supervised and contrastive self-supervised learning. During training, graph augmentation strategies are employed in auxiliary networks to harmonize supervised learning with contrastive learning, enabling the model to utilize unlabeled data effectively. The architecture processes molecular graphs of the POI, E3 ligase, and PROTAC through specialized encoders. Protein graphs are encoded using an edge-weighted GCN (EW-GCN), whereas PROTAC graphs utilize a Convolution with Edge-Node Switching (CensNet) encoder. The encoded representations are then passed to a decoder comprising a modal interaction module and a classifier to produce final predictions. AiPROTAC demonstrated impressive performance across multiple benchmarks. On the PROTAC-DB 2.0 dataset^35^, it achieved an average accuracy of 85.02%, an area under the receiver operating characteristic curve (AUROC) of 0.9192, and an area under the precision-recall curve (AUPR) of 0.8278. We also curated an external test set called PROTAC-ZL by incorporating data not previously disclosed in PROTAC-DB 2.0. On this external test set, the model maintained robust results, with an accuracy of 79.82%, an AUROC of 0.8156, and an AUPR of 0.7504. Further analysis of structurally related samples underscored AiPROTAC’s sensitivity and reliability for PROTAC degradation prediction.

Beyond in silico validation, we further applied AiPROTAC in an AI-driven drug discovery framework, as depicted in Fig. 1b, to incorporate AiPROTAC into PROTAC design pipelines. As a proof of concept, we designed a series of PROTACs targeting androgen receptor (AR) by recruiting cereblon (CRBN) and conducted computational screening. Considering synthetic feasibility, 19 AR-degrading PROTACs were synthesized and experimentally tested. By comparing the experimental results with our AiPROTAC’s predictions, the model correctly identified 14 out of these 19 PROTACs, achieving a prediction accuracy of 74%. Among these 14 PROTACS, five exhibited good degradation capacities (i.e., true positives), while the remaining PROTACs showed poor degradation capacities (i.e., true negatives). Hence, AiPROTAC enables efficient candidate filtering by eliminating low-performing PROTACs, retaining only true positives for synthesis and validation. This significantly accelerates the PROTAC-based drug discovery process by reducing experimental costs and time. Importantly, the screening also yielded a lead AR degrader with drug-like characteristics validated in vitro for its degradation performance. These findings highlight AiPROTAC’s effectiveness and the transformative potential of AI-driven drug discovery in advancing pharmaceutical innovation.

## 2 Results

### 2.1 AiPROTAC framework

AiPROTAC integrates both supervised learning and contrastive learning, as depicted in Fig. 1c. The core of AiPROTAC is the supervised network, which handles the binary classification task, while two auxiliary contrastive learning networks, dedicated to protein and PROTAC tasks, function as branch networks. Notably, these contrastive networks are active only during training, where they leverage both limited labeled data and abundant unlabeled data. The overall training workflow of AiPROTAC can be summarized as follows: batches of molecular graphs representing POI, E3 ligase, and PROTAC are processed in parallel by the classification network and the two auxiliary networks. These networks iteratively update the model parameters until convergence, driven by the optimization objective.

As illustrated in Fig. 1c, in the classification network, protein (POI and E3 ligase) and PROTAC graphs are processed by the EW-GCN encoder and CensNet encoder, respectively, to extract graph-based features. These features are then passed into two parallel attention backbones, each with three input channels: query, key, and value. In the first attention backbone, POI graph features are used as the query, while PROTAC graph features act as both the key and value to derive POI-PROTAC representations. Similarly, in the second attention backbone, E3 ligase graph features serve as the query, and PROTAC graph features function as both the key and value to yield E3 ligase-PROTAC representations. Both attention backbones employ cross-attention to model the PROTAC-induced proximity between POI and E3 ligase. Next, the POI-PROTAC and E3 ligase-PROTAC representations are fed into another parallel attention backbones to obtain POI-viewed and E3 ligase-viewed ternary features. Specifically, when E3 ligase-PROTAC representations serve as the query, and POI-PROTAC representations are used as both key and value, POI-viewed features are derived, and vice versa for the E3 ligase-viewed features. These cross-attention backbones enable the interaction between these two types of representations within the attention space. The resulting POI-viewed and E3 ligase-viewed ternary features are then linearly projected and concatenated, after which they are passed through a multi-layer perceptron to produce the final classification output. In the auxiliary networks, each molecular graph undergoes a graph augmentation process to generate two distinct subgraphs. These subgraphs are subsequently input into a graph neural network (GNN) encoder, which shares the same architecture as the one in the classification network. The contrastive losses for the auxiliary networks are computed based on the GNN encoder outputs, adhering to contrastive learning principles. Finally, the optimization goal *L*_*all*_ consists of three parts: *L*_*sup*_, *L* _*con−p*_ and *L*_*con−c*_, as shown in Fig. 1c. Once trained, AiPROTAC can be used to predict the degradation capacity of a PROTAC, given a set of input molecular graphs (POI, E3 ligase, and PROTAC).

### 2.2 Evaluation of model performance

#### 2.2.1 Data partitioning, parameter settings, and metrics

This research harnessed two datasets, PROTAC-DB 2.0^35^ and PROTAC-ZL, to rigorously evaluate the classification performance of the model. To ensure a fair and reproducible comparison, the data partitioning strategy outlined by Li *et al*.^41^ was followed, with labeled samples in the PROTAC-DB 2.0 dataset initially divided into training, validation, and test sets at an 8:1:1 ratio for hyperparameter optimization. After determining the optimal parameters (Supplementary Table 1), the labeled samples in the PROTAC-DB 2.0 dataset were randomly repartitioned into training and test sets in an 8:2 ratio for subsequent experiments. This repartitioning was repeated across five independent runs with varying random seeds to ensure robustness. The external test set PROTAC-ZL, which contains no overlapping PROTACs with the PROTAC-DB 2.0 dataset, was used to further examine the classification performance of the model. Accuracy, AUROC, and AUPR were adopted as the primary evaluation metrics, while precision, recall, and F1-score were also reported to provide a more comprehensive assessment of the model.

#### 2.2.2 Performance on benchmark datasets

As previously mentioned, we split the labeled samples in the PROTAC-DB 2.0 dataset into training and test sets in an 8:2 ratio, repeating this process five times to create five distinct training and test sets. The model was trained with the optimal parameters on each training set, and thereafter, the trained model was employed to predict the degradation capacities of PROTACs in the corresponding test set. This procedure was repeated across all five sets. To evaluate model performance, we calculated accuracy, precision, recall, F1-score, AUROC, and AUPR based on the predicted and true degradation capacities of PROTACs in the PROTAC-DB 2.0 test sets. Detailed results for all five test sets are provided in Supplementary File 1. Additionally, a separate model was trained on the entire PROTAC-DB 2.0 dataset with optimal parameters, and its performance was assessed on the PROTAC-ZL dataset with accuracy, AUROC, and AUPR as key metrics.

Herein, we implemented ten baseline methods (*i.e*., RF-MACCS, RF-Morgan, SVM-MACCS, SVM-Morgan, DeepPROTACs^41^, DeepPROTACs-Sinput, AiPROTAC-GCN, AiPROTAC-GAT, AiPROTAC-Rattention, and AiPROTAC-RgraphCL), and the corresponding implementation code is publicly accessible at https://github.com/LiZhang30/AiPROTAC. Among these, RF-MACCS, RF-Morgan, SVM-MACCS, and SVM-Morgan are classical machine learning (ML) models, underpinned by two fundamental algorithms: Random Forest (RF) and Support Vector Machine (SVM). In these models, proteins are characterized by amino acid residue sequences, while ligands are encoded using molecular access system (MACCS) keys or Morgan fingerprints. DeepPROTACs^41^, one of the few models specifically designed for predicting the degradation capacities of PROTACs, serves as a reference. The model DeepPROTACs-Sinput, which shares a similar network structure with DeepPROTACs, utilizes the same input format as AiPROTAC. Moreover, AiPROTAC-GCN, AiPROTAC-GAT, AiPROTAC-Rattention, and AiPROTAC-RgraphCL are variants of the AiPROTAC framework. The first two variants substitute the GNN encoder in AiPROTAC with the graph convolutional network (GCN)^42^ and graph attention network (GAT)^43^, respectively. The latter two variants are modified by removing all attention backbones and contrastive learning networks from AiPROTAC.

As summarized in Figs. 2a and 2b, AiPROTAC achieves average accuracy, precision, recall, f1-score, AUROC, and AUPR values of 0.8502, 0.8298, 0.8467, 0.7935, 0.9192, and 0.8278, respectively, on the test set of the PROTAC-DB 2.0 dataset. The corresponding standard deviations (SD) for these metrics are 0.0184, 0.0494, 0.0450, 0.0194, 0.0097, and 0.0505. Figs. 2c and 2d present the results of all baselines on the six metrics in the PROTAC-DB 2.0 test set with the random seed of 1, while the results of the other four experiments are shown in Supplementary Fig. 1. Fig. 2e exhibits the accuracy (0.7928), AUROC (0.8156), and AUPR (0.7504) for AiPROTAC on the PROTAC-ZL dataset. Compared to the ten baseline methods described earlier, AiPROTAC demonstrates superior performance on both the PROTAC-DB 2.0 and PROTAC-ZL datasets (Fig. 2 and Supplementary Tables 2 and 3). Furthermore, the low SD across all metrics in the PROTAC-DB 2.0 dataset highlights the robust stability of AiPROTAC. Notably, the average value (0.7862) of the key metrics (*i.e*., accuracy, AUROC, and AUPR) for AiPROTAC on the PROTAC-ZL dataset is approximately 25% higher than that of DeepPROTACs^41^ (0.5450), emphasizing AiPROTAC’s excellent generalization performance. Moreover, AiPROTAC consistently outperforms its four variants across all metrics on both benchmark datasets, showcasing the effectiveness of GNN encoders, attention-based modal interactions, and contrastive learning networks in this study.

**Fig. 2.**
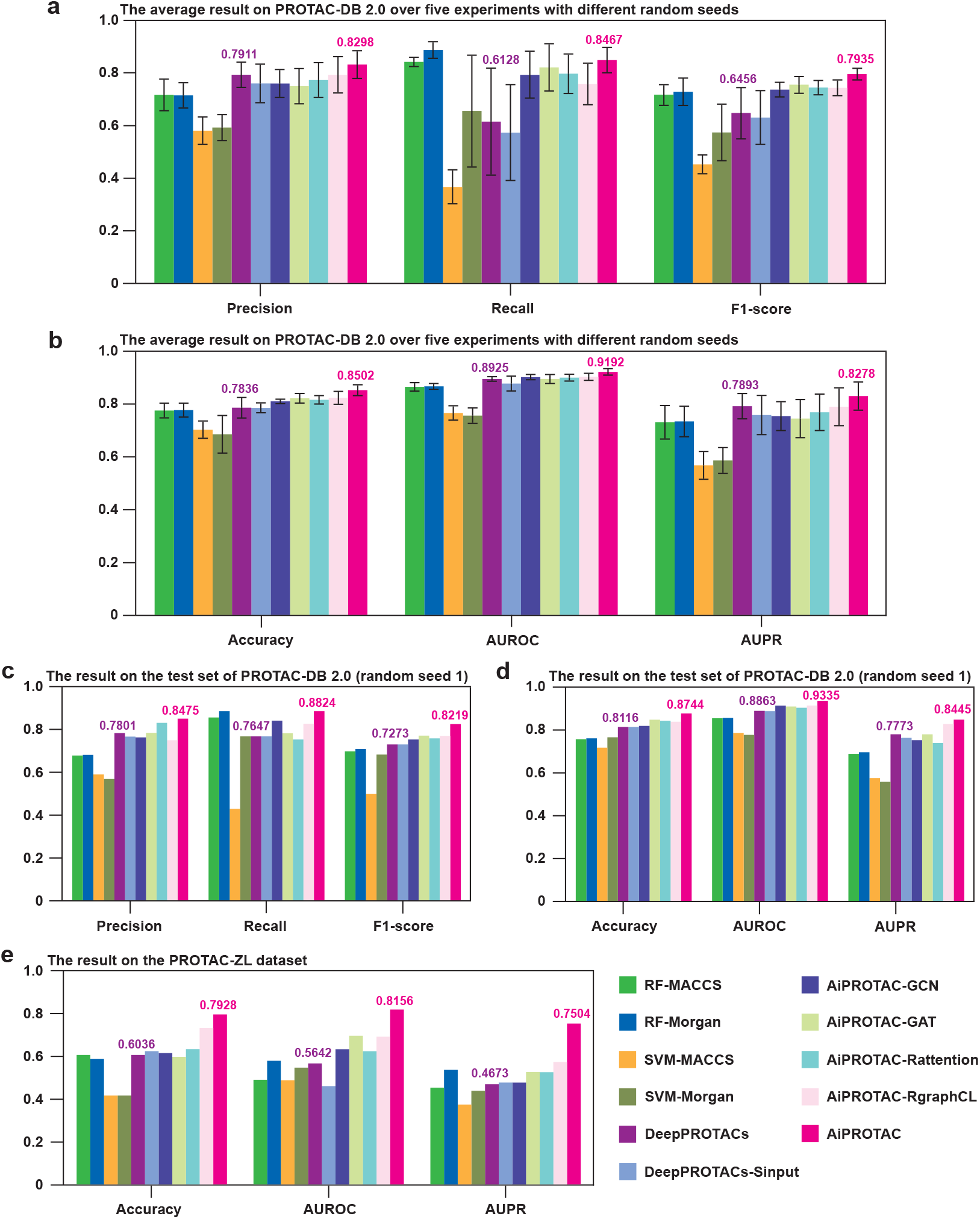
Prediction results on the public and extended datasets. **(a-b)**. The average result of five experiments with different random seeds on the PROTAC-DB 2.0 dataset using eleven methods. **(c-d)**. The experimental result of eleven methods on the PROTAC-DB 2.0 dataset with random seed 1. **(e)**. The experimental result of eleven methods on the PROTAC-ZL dataset. The average value (0.7862) of accuracy, AUROC and AUPR for AiPROTAC is approximately 25% higher than that (0.545) of DeepPROTACs.

### 2.3 Case studies on model sensitivity

Two case studies were conducted to evaluate the sensitivity of AiPROTAC to various inputs, utilizing data sourced from separate investigations^44, 45^. In particular, fourteen samples related to histone deacetylases (HDACs), two series of PROTACs, and CRBN were extracted from a recent study^44^. In this study, the first series of PROTACs (A1-6) incorporates a vorinostat-like HDAC ligand linked by an alkyl chain, whereas the second series (B1-4) features a selective benzimidazole-based HDAC6 ligand^46^. Similarly, ten samples involving five POIs (*i.e*., FER, AAK1, GAK, CHK2, and IRAK1), two PROTACs, and CRBN were collected from the study^45^. All biomolecules were converted into molecular graphs, and each sample was labeled according to in vitro experimental data, including half maximal degradation concentration (*DC*_*50*_), maximal degradation (*D*_*max*_), and Western blotting results, as provided in the aforementioned studies. The trained AiPROTAC model was then applied to predict the labels of all samples.

Drawing from both experimental and predicted degradation data for proteins targeted by CRBN-recruiting PROTACs (Fig. 3), we accessed AiPROTAC’s prediction accuracy, achieving 100% and 70% across two distinct scenarios, respectively. In the first scenario, when any of HDAC1, HDAC4, or HDAC6 was designated as the POI in the sample, the entire sample’s input variations originated solely from differences in the PROTAC’s linkers or target ligands. Under this scenario, AiPROTAC maintained flawless prediction accuracy. In the second scenario, when either SIAIS352008 or SIAIS262039 was used as the PROTAC input in the sample, the entire sample’s input variations were exclusively due to differences in POIs, and the model still attained a robust 70% accuracy. These findings demonstrate that AiPROTAC exhibits robust sensitivity to variations in model inputs, whether stemming from small molecule modifications or differences in target proteins, thereby reinforcing its reliability and efficacy.

**Fig. 3.**
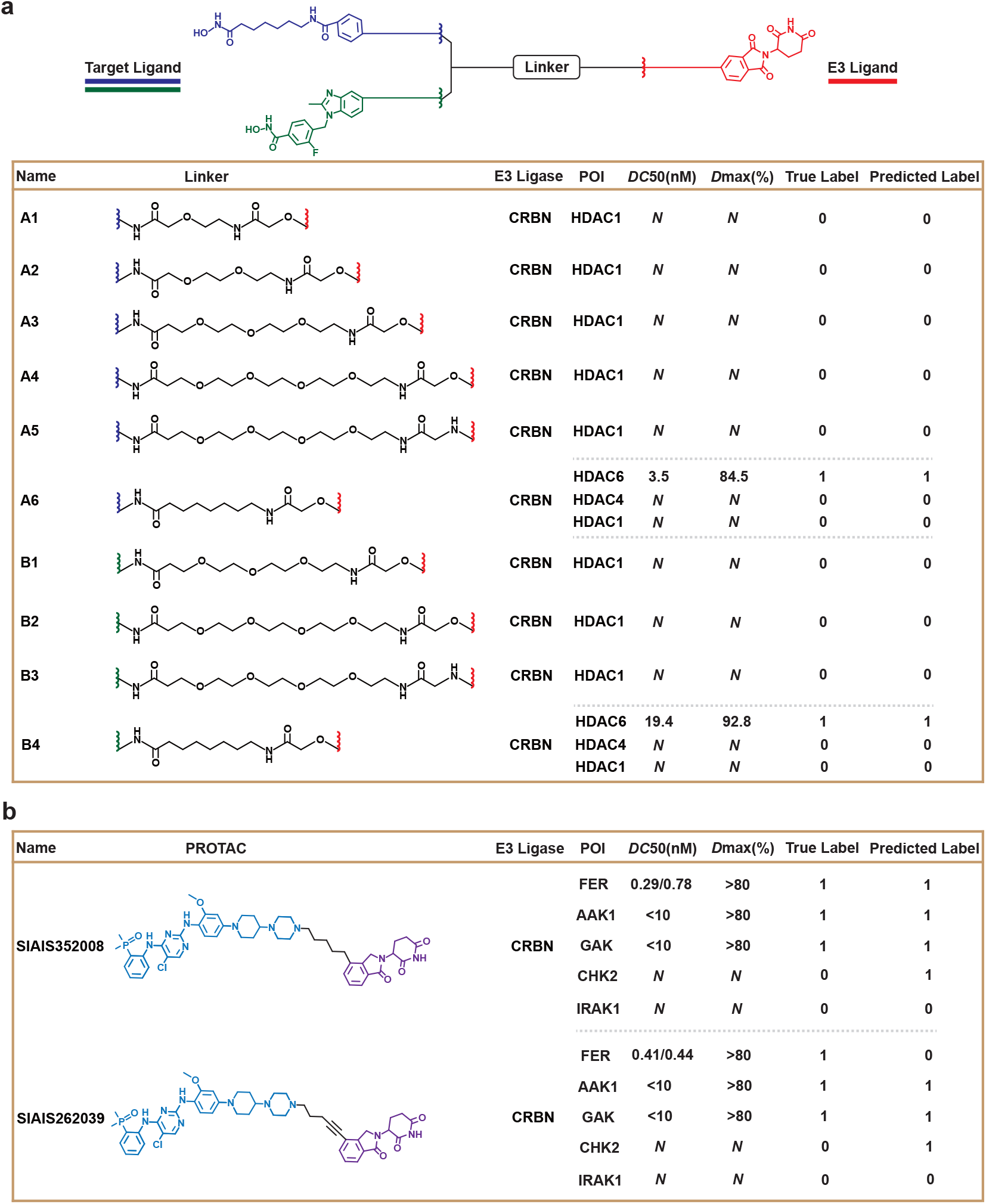
Experimental results for evaluating the sensitivity of AiPROTAC. **(a)**, Experimental and predicted degradation results of HDACs targeted by two series of PROTACs recruiting CRBN. **(b)**, Experimental and predicted degradation results of five different proteins targeted by two PROTACs recruiting CRBN. Note: The symbol *N* means *DC* 50>100 nM or *D* max<80% according to the results of Western blotting.

### 2.4 AI-assisted discovery of AR degrader

#### 2.4.1 AI-based drug development paradigm

The typical drug development paradigm begins with chemists designing numerous drug-like molecules that target specific therapeutic sites. This is followed by chemical synthesis and a series of systematic in vitro experiments to assess their efficacy and properties^47–49^. In contrast, the approach emphasized in this study introduces an advanced paradigm, wherein an AI-driven predictor is leveraged prior to the synthesis of expert-curated molecules. The role of the AI predictor is to preemptively eliminate compounds considered undruggable (Fig. 1b). This cutting-edge strategy not only enhances the efficiency of identifying viable drug candidates but also offers substantial benefits in terms of cost reduction, accelerated development timelines, and improved success rates throughout the drug discovery process.

#### 2.4.2 Discovery process of lead candidate PROTAC

To advance the practical application of AiPROTAC, evaluate the efficacy of AI-driven drug development paradigms in real-world settings, and identify additional lead PROTAC candidates, we initiated a project focused on the development of AR degraders. This comprehensive effort encompassed systematic research, including the analysis of pathological and pharmacological mechanisms, molecular design, virtual screening, chemical synthesis, and in vitro experiments. The AR, also known as NR3C4 (*i.e*., nuclear receptor subfamily 3, group C, member 4), is an androgen-dependent transcription factor featuring four functional domains: the ligand-binding domain, DNA-binding domain, hinge region, and N-terminal domain^50^. Degradation of the AR protein holds significant promise for overcoming resistance mechanisms associated with current AR-targeted therapies, thereby offering a novel therapeutic strategy for patients diagnosed with prostate cancer^51–53^.

Fig. 4a illustrates the design of two groups of AR-targeting PROTACs, comprising a total of 18 compounds. The first group consists of 12 PROTACs (GT01-12), developed using nicotinamide derivative (analog of ARV-110 warhead) as the target protein ligand, S-substituted Lenalidomide (a novel CRBN ligand developed by Yang group) as the E3 ligase ligand, and various linear alkyl chains of differing lengths as linkers. The second group includes 6 PROTACs (GT13-18), which also utilize the same CRBN ligand but differ by employing Enzalutamide as the warhead. We employed AiPROTAC to predict the degradation capacities of these AR-targeting PROTACs, followed by the synthesis of the compounds and evaluation of their ability to induce AR degradation in the LNCaP prostate cancer cell line. The predicted results were highly consistent with the in vitro experimental outcomes (Fig. 4b), with 8 out of 12 PROTACs in the first group and 5 out of 6 PROTACs in the second group being accurately predicted. This high degree of alignment underscores the effectiveness of AiPROTAC in forecasting the degradation potential of PROTACs targeting the AR protein. However, based on assay results, only GT02, GT03, GT05, and GT09 among the 18 PROTACs met the criteria for good degradation. To develop higher-quality degraders, we extended the design of the first group by adjusting the binding position of the linker on the benzene ring in the CRBN ligand and designed GT19. AiPROTAC recognized GT19 as possessing strong AR-targeting degradation capacity. Western blotting confirmed that GT19 indeed exhibits a higher level of AR degradation.

**Fig. 4.**
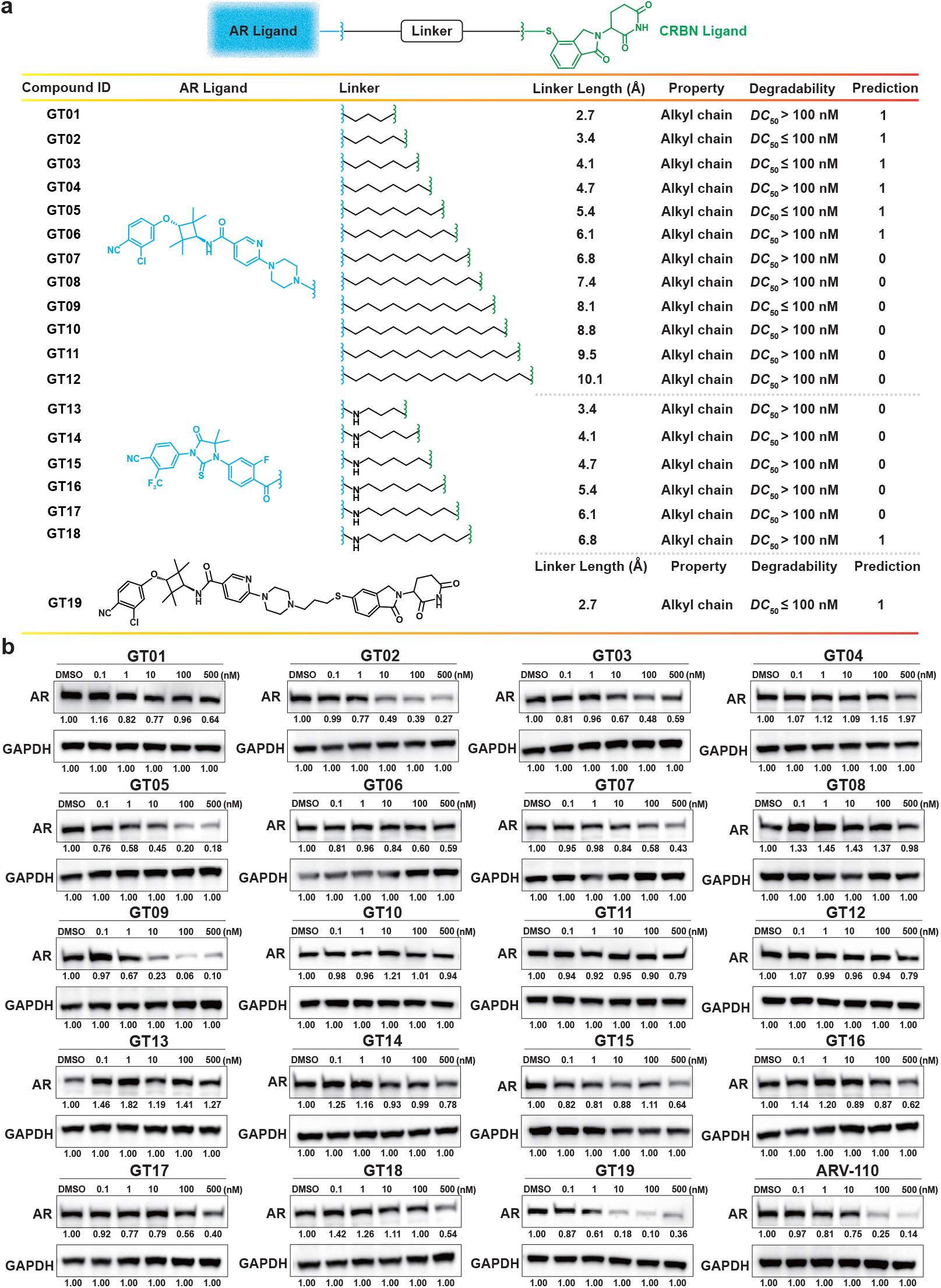
The synthesis of PROTACs and in vitro experiments. **(a)**, GT01-12 and GT19 utilize a nicotinamide derivative as the AR ligand (with GT19 binding to CRBN differently from GT01-18), while GT13-18 use Enzalutamide. **(b)**, Western blotting analysis and densitometric quantification of AR protein in LNCaP cell lines treated with compounds at 0.1, 1, 10, 100, and 500 nM for 24 h. **22/24**

In our analysis of 19 PROTACs using Western blotting, ARV-110 served as the control. Among these, GT09 and GT19 demonstrated superior AR degradation, overall outperforming ARV-110 at concentrations below 100 nM within a 24-hour treatment period (Fig. 4b). Notably, the linker in GT19 is significantly shorter than that in GT09, suggesting that GT19 may have better bioavailability and druggability^54^. In light of this, we prioritized GT19 for further studies. To validate the degradation mechanism of GT19, we synthesized the negative control compound GT19NC by methylating phthalimide, the key part that binds to CRBN. We first examined the CRBN binding affinity of GT19 and GT19NC utilizing the HTRF human cereblon binding kit (Supplementary Fig. 2a). The *IC*_*50*_ values for GT19 was measured to be 0.15 µM, significantly stronger than the control compounds Lenalidomide and Pomalidomide, indicating a robust binding affinity between GT19 and CRBN. In contrast, GT19NC exhibited negligible binding to CRBN, with an *IC*_*50*_ value exceeding 10 µM. Furthermore, Western blot showed that AR was obviously degraded by GT19 at the concentration of 100 nM, while GT19NC failed to degrade AR (Supplementary Fig. 2b). These results confirm that AR PROTACs recruit and bind to CRBN ligase, thereby facilitating AR degradation through the ubiquitin-proteasome pathway. Consequently, GT19 is identified as the lead candidate PROTAC in this study, with potential for further development into a clinically applicable AR degrader. In conclusion, AiPROTAC successfully predicted the degradation labels of 14 out of 19 PROTACs (Supplementary Tables 4 and 5), achieving a 74% prediction accuracy rate. This demonstrates its excellent predictive performance in practical application scenarios. Moreover, this study serves as a compelling case example of utilizing AiPROTAC in drug development, effectively identifying and validating the lead PROTAC GT19 targeting AR, which can be further developed into clinically available drugs. The AR degrader development project exemplifies the AI-driven drug development paradigm, providing an example of using AiPROTAC.

## 3 Discussion

We present AiPROTAC, an innovative model designed to accelerate the development of potent PROTACs by integrating graph augmentation, two advanced graph encoders, cross-attention, and multiple perceptrons within a unified, learnable framework that supports both supervised and self-supervised learning. AiPROTAC excels at virtually screening candidate degraders by predicting the potential of a given PROTAC to induce the degradation of a POI through the recruitment of a corresponding E3 ligase. This task of predicting PROTAC-induced degradation is pivotal for successful degrader development^55^. However, the lack of computational tools tailored to this task was a key driver in the creation of AiPROTAC. To address the challenge of limited sample size in PROTAC-induced degradation prediction, we pursued two strategies. First, we manually curated labeled samples to expand the existing dataset. Second, we constructed two auxiliary tasks to capitalize on unlabeled samples, employing contrastive learning to strengthen the predictive capability of the model. Furthermore, we developed two specialized encoders, EW-GCN and CensNet, to more effectively characterize proteins and PROTACs, thus addressing the underutilization of physicochemical property information in molecular graph edges. The EW-GCN encoder is derived by transforming the standard GCN into an EW-GCN network with a residual connection, whereas the CensNet encoder is obtained by upgrading the original CensNet to its residual form. Additionally, we introduced a two-stage cross-attention mechanism to emulate the logic of wet-lab experiments. In the first stage, cross-attention models the proximity between the POI and E3 ligase induced by the PROTAC, while in the second stage, it captures features of the ternary complex from two complementary perspectives.

AiPROTAC demonstrates an impressive accuracy of 85.02% in predicting sample labels within the PROTAC-DB 2.0 test set, outperforming all baseline methods. It also ranks as the top performer in two critical metrics, AUROC and AUPR (0.9192 and 0.8278). When applied to the hand-crafted PROTAC-ZL test set, AiPROTAC achieves the highest accuracy, AUROC, and AUPR (79.82%, 0.8156, and 0.7504) compared to state-of-the-art models. Notably, in two specific cases warranting further attention, AiPROTAC attains perfect accuracy (100%) and a commendable 70% accuracy in predicting PROTAC-induced degradation, underscoring its capacity to differentiate molecular types and discern structural nuances of compounds. In one instance, the model’s input variation stems from the PROTACs, while in the other, it is driven by the target proteins. The AI-driven degrader development system proposed in this paper marks a significant evolution beyond traditional biological experimentation. This system has already been leveraged to identify AR degraders, offering promising treatment alternatives for the 1.4 million cases of prostate cancer diagnosed globally each year^56^. More specifically, by applying the AI-assisted degrader development process outlined above, a lead candidate molecule (GT19) was identified, which induces AR protein degradation more effectively than ARV-110. This application reveals the potential of AiPROTAC in reducing costs, accelerating development timelines, and improving success rates throughout the drug development pipeline.

While AiPROTAC has demonstrated notable success in several areas, there remains significant potential for further enhancement in the long term. As a data-driven approach, even with the incorporation of unlabeled data via an unsupervised learning framework, AiPROTAC faces inherent performance limitations due to the constraints of available data. Therefore, the continued collection of additional data is both necessary and urgent. Furthermore, by harnessing large-scale datasets of small molecules and protein sequences from public databases, a pre-trained large chemical model^57^ can be employed as an encoder to capture fine-grained features of input molecules, thereby enhancing the accuracy of degradation predictions. The ternary complexes induced by PROTACs are critical for targeted protein degradation, making the spatial characterization of these complexes and their binding domains a key area of focus. Drawing inspiration from recent studies^58, 59^, all-atom modeling offers a promising approach to address this challenge. Additionally, transforming the binary classification task into a regression task by predicting values such as *DC*_*50*_ and *D*_*max*_ could provide a more direct assessment of the degradation efficacy of PROTACs on POIs. However, given the current rate of data generation and publication in this field, the available data in the short term may not suffice to support such regression tasks. An alternative approach is to frame the task as a multi-class classification problem, incorporating additional chemical indicators alongside *DC*_*50*_ and *D*_*max*_. This would expand the class labels beyond two, offering a more nuanced representation of the actual degradation levels of PROTACs. In summary, AiPROTAC marks a pioneering effort in applying AI to the PROTAC domain, underscoring the transformative potential of AI in the degrader development pipeline and laying the foundation for an AI-driven drug discovery paradigm.

## 4 Methods

### 4.1 Datasets

We examined the performance of prediction models on account of PROTAC-DB 2.0 and PROTAC-ZL datasets. The former comes from a web-accessible database published by Hou *et al*.^35^, while the latter is manually collected from recently published researches^44, 45, 60–68^. The samples in the original PROTAC-DB 2.0 database are not labeled, and the developers of DeepPROTACs^41^ have not made their data public. Unlike previous situations, the mentioned two datasets we released at https://github.com/LiZhang30/AiPROTAC can be utilized for developing prediction models without preprocessing. A pair of quantitative indicators, *DC*_*50*_ and *D*_*max*_, are exploited to label our data according to the criterion: compounds with *DC*_*50*_ lower than 100 nM and *D*_*max*_ higher than 80% are marked as good degraders; otherwise, they are labeled as bad degraders. As illustrated in Fig. 5a, the labeled data in PROTAC-DB 2.0 dataset was first split into training, validation, and test sets at an 8:1:1 ratio to optimize the model’s hyperparameters. After optimization, the labeled data was re-split into training and test sets at an 8:2 ratio for subsequent experiments, reducing variability across training trials. Performance was evaluated on the 20% test set and further validated on the PROTAC-ZL dataset employing the same metrics.

**Fig. 5.**
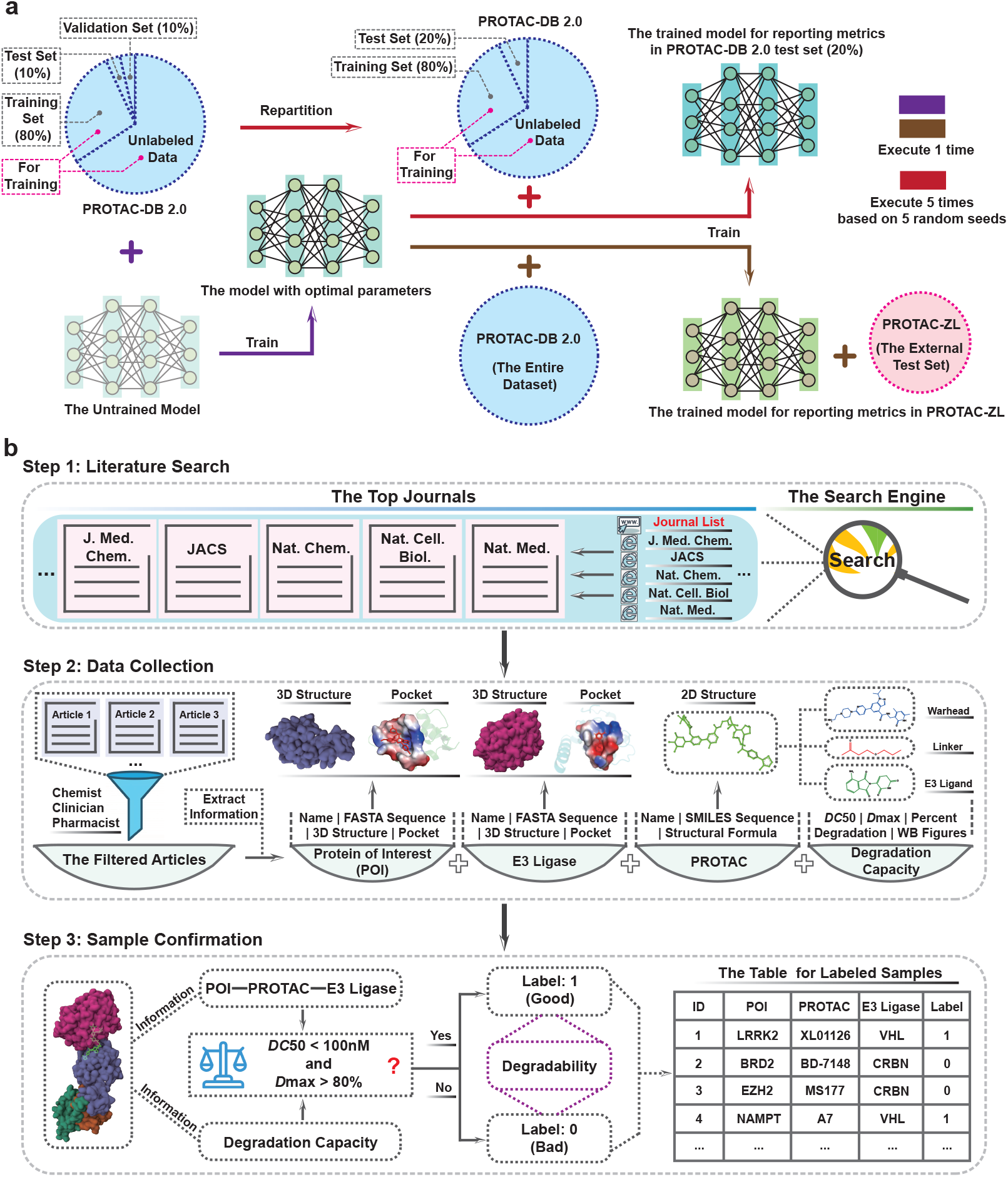
The utilization and expansion of data in this study. **(a)**, The process of training and evaluating the model using our two datasets. Optimal parameters of the model are firstly determined using the PROTAC-DB 2.0 dataset. After training the model with the repartitioned training set and unlabeled data in the PROTAC-DB 2.0 dataset, the metrics are reported on the PROTAC-DB 2.0 test set to evaluate the model. The same metrics are also reported on the PROTAC-ZL dataset to further assess the model trained with the entire PROTAC-DB 2.0 dataset. **(b)**, The construction process of the PROTAC-ZL dataset. The implementation steps include literature search, data collection and sample confirmation.

In this research, after a series of preprocessing steps on the raw data, we obtained the PROTAC-DB 2.0 dataset used herein. It contains 5,388 samples, consisting of 1,036 labeled samples and 4,352 unlabeled samples. Fig. 5b depicts three steps involved in constructing the PROTAC-ZL dataset: literature search, data collection and sample confirmation. We initially acquired numerous literatures regarding PROTACs and filtered out the invalid ones. After removing these literatures already included in the PROTAC-DB 2.0 database, we gathered all necessary information for conducting experiments from remaining valid literatures. Based on the collected wet lab results (*i.e*., *DC*_*50*_, *D*_*max*_, percent degradation and Western blotting), all samples were labeled and their complete information was recorded. The PROTAC-ZL dataset contains 111 labeled samples, involving 76 PROTACs not reported in the PROTAC-DB 2.0 dataset. Each PROTAC in the PROTAC-ZL dataset has been split into three parts: warhead, linker, and E3 ligand. More detailed information related to the two datasets can be found in Supplementary Tables 6 and 7.

### 4.2 Model input representation

The input for AiPROTAC contains three types of molecular graphs: the POI graph, the PROTAC graph and the E3 ligase graph. POI and E3 ligase graphs are generated with the same manner in this study. As displayed in Fig. 6a, the enlarged section of the protein structure shows the actual biological significance of nodes and edges in the protein graph. Concretely, each residue is regarded as a node and we add an edge between two nodes if the distance between the corresponding residues is less than 8 Å. In addition, the physical and chemical properties of residues are applied to construct the node feature, while the distance and similarity between two residues are employed to build the edge feature. The magnified portion of the molecule conformation illustrates the corresponding elements in a molecule represented by nodes and edges in the molecule graph. Specifically, each atom is regarded as a node and we add an edge between two nodes if there is a chemical bond between the corresponding atoms. Besides, the node and edge feature are derived from the properties of atoms and chemical bonds, respectively.

**Fig. 6.**
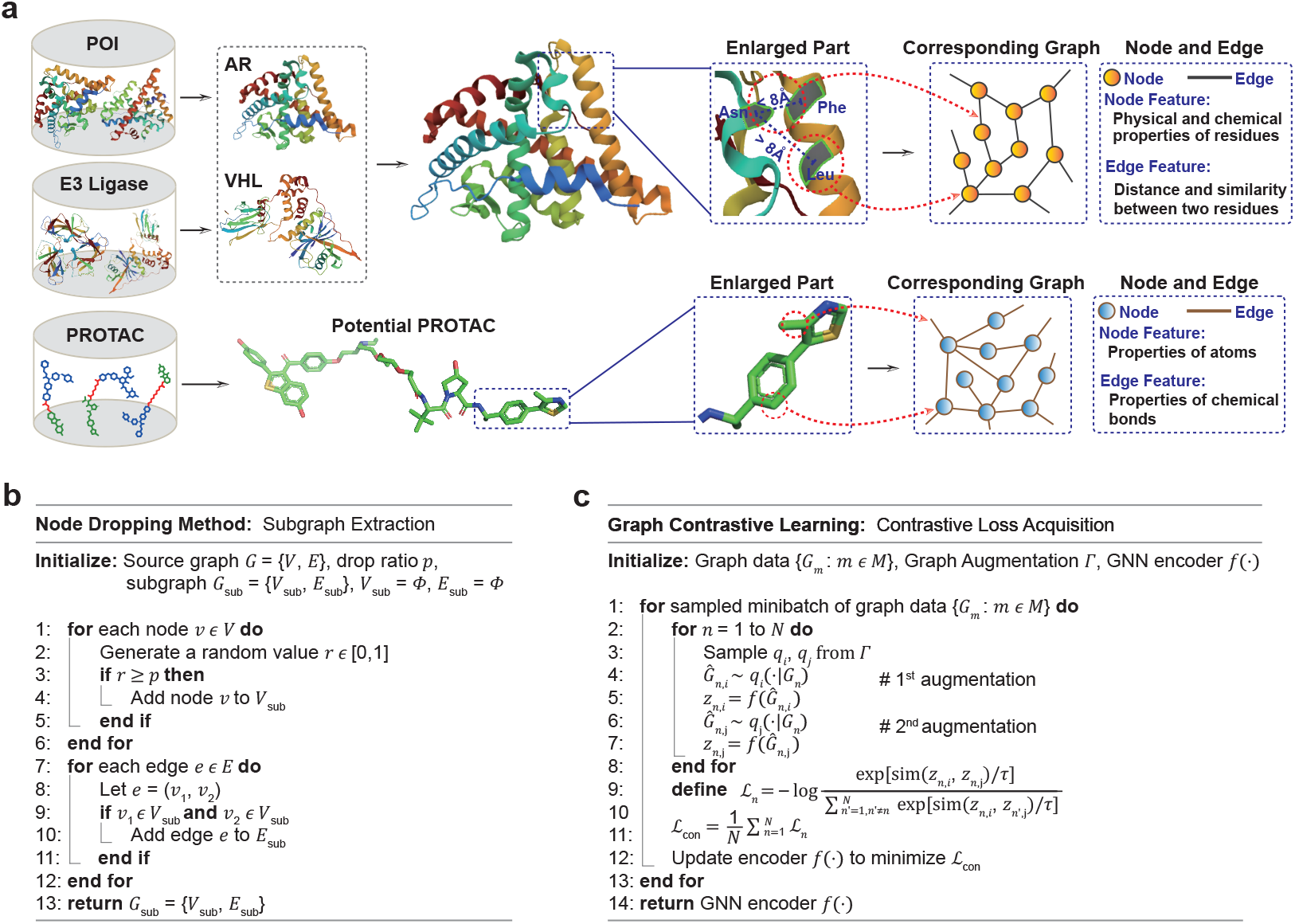
Molecular graph construction and graph contrastive learning. **(a)**, Generating a molecular graph. For a protein, each residue is a node, and edges are formed if the distance between residues is less than 8 Å. Node features are defined by residue properties, while edge features are determined by residue distance and similarity. For a PROTAC, each atom is a node, with edges representing chemical bonds. Atomic and bond properties define node and edge features, respectively. **(b)**, Graph augmentation principle. Two distinct subgraphs are generated from a molecular graph using the node dropout method. **(c)**, Graph contrastive learning process. Graph contrastive learning involves applying graph augmentation to create positive pairs, encoding graphs with a GNN, and optimizing a contrastive loss function.

Herein, the POI graph, the PROTAC graph and the E3 ligase graph are defined as *G*_*a*_ = (*V*_*a*_, *E*_*a*_), *G*_*b*_ = (*V*_*b*_, *E*_*b*_) and *G*_*c*_ = (*V*_*c*_, *E*_*c*_), where |*V*| is the set of nodes and |*E*| is the set of edges. *m* = |*V*| and *n* = |*E*| denote the number of nodes and edges, respectively. *v*_*i*_ *∈V* is the *i*-th node and *e*_*ij*_ *∈ E* is the edge between *i*-th and *j*-th nodes, where *i, j ∈{*1, 2, …, *m}* . ***A***_*e*_ *∈*ℝ^*m×m*^ signifies the adjacency matrix of a graph, where each element ***A***(*i, j*) denotes whether there is an edge between node *i* and node *j*. 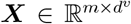 denotes the node feature matrix, where each node is associated with a *d*^*v*^-dimensional feature vector ***x***. 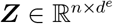 signifies the edge feature matrix, where each edge has a *d*^*e*^-dimensional feature vector ***z*** (Supplementary Tables 8 and 9). ***A***_*e*_*∈* ℝ^*n×n*^ denotes the adjacency matrix of a corresponding line graph, and ***A***_*e*_(*s, t*) equals to 1 if edge *s* and edge *t* are connected by a node in the original graph, otherwise 0.

### 4.3 Implementation details of AiPROTAC

#### 4.3.1 Overview of AiPROTAC’s components

To predict the degradation capacity of PROTACs with known target proteins and E3 ligases, we have proposed a framework that integrates supervised and contrastive learning. This approach incorporates the graph augmentation mechanism and the GNN encoder for feature enhancement, along with the attention backbone and the multiple perceptrons for feature fusion.

#### 4.3.2 Graph augmentation mechanism

In this work, the graph augmentation mechanism^69^, which samples two distinct subgraphs from a molecular graph, serves as the basis for implementing graph contrastive learning and utilizing unlabeled samples. Supplementary Fig. 3a illustrates the process of extracting two different subgraphs from the source graph by randomly discarding 20% of the nodes, using two separate random seeds. Furthermore, the specific logic of the node dropping method is pictured in Fig. 6b. First, a subset of nodes from the source graph is discarded according to the given drop ratio. Then, each edge in the source graph is examined, and it is included in the subgraph only if both of its connected nodes are retained (*i.e*., neither is in the discarded node set).

#### 4.3.3 GNN encoder

We designed the EW-GCN and CensNet encoders to extract features from molecular graphs. The EW-GCN encoder is specifically tailored to process POI and E3 ligase graphs, with its network structure illustrated in Fig. 1d. EW-GCN operates as a graph encoder, stacking multiple EW-GCN layers in sequence with a residual mechanism strategy. Particularly for protein graphs, the EW-GCN layer effectively leverages the edge weight information to facilitate more efficient message passing

between nodes. The inputs to the EW-GCN layer are the node adjacency matrix, the node feature matrix and the edge feature matrix. In protein graphs, the edge feature, including the distance between two residues and the cosine similarity of two residue features, are employed to calculate the weight on each edge. Let the edge weight matrix be *A* ^*p*^ *∈*ℝ^*m×m*^, then, the Laplacianized edge-weighted node adjacency matrix is given by:

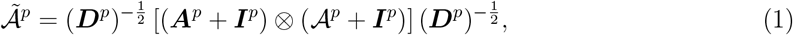

where ***D***^*p*^ represents the diagonal degree matrix of ***A***^*p*^ + ***I***^*p*^, and ***I***^*p*^ is the identity matrix. The operator ⊗ denotes the Hadamard product. The layer-wise propagation rule for node features in the (*l* + 1)-th layer is mathematically expressed as:

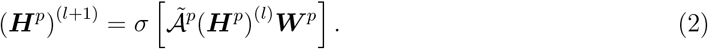

Herein, *σ* is the ReLU activation function, ***H***^*p*^ states the node feature matrix at the *l*-th layer, and ***W*** ^*p*^ is a layer-specific trainable weight matrix. At the final layer of the EW-GCN encoder, we aggregate the features of all nodes in the protein graph through the element-wise sum, resulting in the *d*-dimensional output vector ***x***^*p*^ .

The CensNet encoder, designed for PROTAC graphs, utilizes the CensNet layer^70^ as its foundational network. Similar in structure to the GCN encoder, it sequentially connects multiple CensNet layers based on the residual mechanism. As depicted in Fig. 1e, each CensNet layer consists of two types of sublayers: a node layer and an edge layer. The inputs to each CensNet layer include a node adjacency matrix and a node feature matrix, as well as an edge adjacency matrix and an edge feature matrix. Unlike the message passing rules in conventional GNNs, the propagation rules in the CensNet layer are characterized by two aspects. In the node layer propagation, all inputs are processed to update node features, while the line graph advances without any modifications. We first define the Laplacianized node adjacency matrix as:

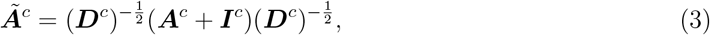

where ***D***^*c*^ signifies the diagonal degree matrix of ***A***^*c*^ + ***I***^*c*^, and ***I***^*c*^ means the identity matrix. Then, the layer-wise propagation rule for node features in the (*l* + 1)-th layer can be regarded as:

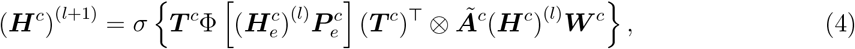

where 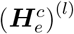 and (***H***^*c*^)^(*l*)^ indicate the edge feature matrix and the node feature matrix at the *l*-th layer, respectively. ***T*** ^*c*^*∈* ℝ^*m×n*^ is a binary transformation matrix, with values that reflect the association between nodes and edges. The operator Φ denotes diagonalization. Moreover, 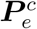 and ***W*** ^*c*^ are defined as learnable weight matrices, used for edge features and node features, respectively. In the edge layer propagation, the updated node feature matrix is combined with the line graph to update the edge feature matrix. Specifically, we calculate the Laplacianized edge adjacency matrix as:

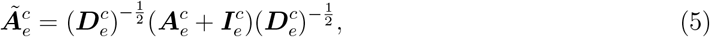

where 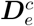 means the diagonal degree matrix of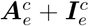, and 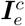is the identity matrix as well.

Analogously, the propagation rule for edge features in the (*l* + 1)-th layer is given by:

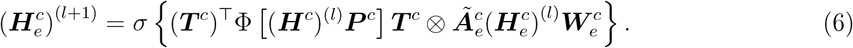

It is clear that ***P*** ^*c*^ and 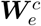 are two additional learnable weight matrices, applied to node and edge features, respectively. Overall, the mechanism of alternately updating node and edge features within a CensNet layer facilitates information exchange between nodes and edges in the graph network, thereby enhancing the representational capacity of the layer. In the CensNet encoder, an element-wise sum is employed to aggregate the features of all nodes in the PROTAC graph, yielding the *d*-dimensional output vector ***x***^*c*^ .

#### 4.3.4 Attention backbone

As exhibited in Supplementary Figs. 3b and 3c, various tricks are adapted in the core scaled dot-product attention mechanism to form the attention backbone, which acts as the basic unit of a two-stage cross-attention network for obtaining POI-viewed and E3 ligase-viewed ternary features. Here, the *i*-th head of the cross-attention, *CH*_*i*_, is first represented as:

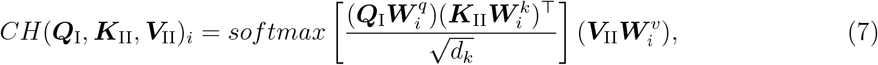

where at the theoretical level, ***Q***_I_, ***K***_II_ and ***V***_II_ correspond to query, key and value, respectively. Besides, ***Q***_I_*≠****K***_II_ = ***V***_II_. The *d*_*k*_ denotes the dimension of each key vector in the ***K***_II_ matrix. 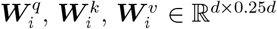 are linear projection matrices. Naturally, we can express the cross multi-head attention as:

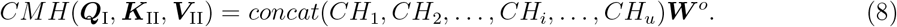

The *u* indicates the number of heads in the cross-attention, and ***W***^*o*^ *∈*ℝ^*d×d*^ is also functioned as a linear projection matrix. In the initial cross-attention stage, the graph vectors 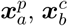 and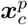, representing POI, PROTAC and E3 ligase are the network inputs to simulate the PROTAC-induced proximity of POI and E3 ligase. In detail, the implementation involves setting the input of the ***Q***_I_ channel to 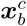 ^*c*^, while the input of the ***K***_II_ = ***V***_II_ channels is set to 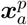 and 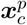 in two different attention backbones, thereby generating POI-PROTAC and E3 ligase-PROTAC representations, respectively. These two representations are then fed into the second cross-attention stage to get the final POI-viewed and E3 ligase-viewed ternary features. The process is similar to the first stage: the ***Q***_I_ channel is set to the E3 ligase-PROTAC representation, and the ***K***_II_ = ***V***_II_ channels are set to the POI-PROTAC representation to produce the *d*-dimensional POI-viewed ternary feature 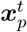 . Conversely, setting the ***Q***_I_ channel to the POI-PROTAC representation and the ***K***_II_ = ***V***_II_ channels to the E3 ligase-PROTAC representation yields the *d*-dimensional E3 ligase-viewed ternary feature 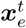 .

#### 4.3.5 Optimization

A linear projection network for optimizing ternary features and a multi-layer perceptron for feature fusion are built with the perceptron as the fundamental layer (Supplementary Figs. 3d and 3e). The forward pass from layer *l* to *l* + 1 in the above two networks is defined as:

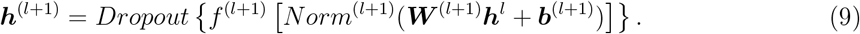

Here, ***h***^(*l*+1)^ means the activations of layer *l* +1, computed by applying a linear transformation to the activations of the previous layer ***h***^*l*^ with the weight matrix 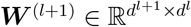 and bias vector 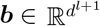. The dimensions *d*^*l*^ and *d*^*l*+1^ represent the number of neurons in layers *l* and *l* + 1, respectively, defining the input and output dimensions for the transformation. The result is then normalized via *Norm*^(*l*+1)^, activated by *f* ^(*l*+1)^, and regularized through dropout to improve generalization.

The considered loss functions comprise supervised loss and contrastive loss. In the supervised task, the binary cross entropy loss *L*_*sup*_ is employed to optimize model predictions for binary classification. It is formulated as:

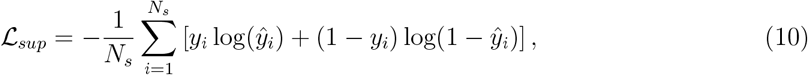

where *N*_*s*_ is the number of samples, *y*_*i*_ ∈{0, 1} denotes the ground truth label for the *i*-th sample, and 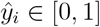 signifies the predicted probability produced by a sigmoid activation function.

Fig. 6c provides a detailed illustration of the working mechanism of graph contrastive learning. During training, after initializing the graph data, graph augmentation and GNN encoder, a minibatch of *N* graphs are randomly sampled and processed through contrastive learning, resulting in 2*N* augmented graphs and the contrastive loss. Specifically, the *n*-th graph *G*_*n*_ in the minibatch undergoes graph augmentation to generate two correlated views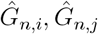, forming a positive pair, where 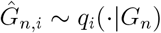 and 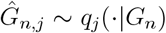. Besides, negative pairs are not explicitly sampled but generated from the other *N* − 1 augmented graphs. The GNN encoder is then used to extract graph-level representation vectors *z*_*n*,*i*_, *z*_*n*,*j*_ from augmented graphs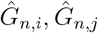. Importantly, we adopt the normalized temperature-scaled cross entropy loss (NT-Xent)^71^ to implement the contrastive loss, and the cosine similarity function is denoted as sim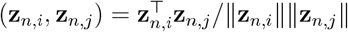. The NT-Xent _*n*_ for the *n*-th graph, and the overall contrastive loss _con_ across all positive pairs are computed as follows:

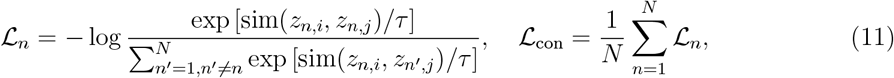

where *τ* denotes the temperature parameter. For auxiliary tasks involving proteins and PROTACs, we define two distinct losses, *L*_*con−p*_ and *L*_*con−c*_. Specifically, *L*_*con−p*_ focuses on distinguishing among various proteins in the graph representation space, while *L*_*con−c*_ captures the distinctions among different PROTACs in that space. Consequently, the final optimization objective is written as:

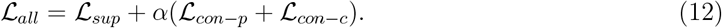

Parameter *α* refers to the loss allocation coefficient, which is set to 0.05 in our experiments.

### 4.4 General experimental methods

#### 4.4.1 Chemical materials

All chemicals were obtained from commercial suppliers, and used without further purification, unless otherwise indicated. Compound 1 were prepared according to the patent WO2019023553. Compound 2, 3 and 4 were prepared according to the patent WO2019196812. HPLC preparation was performed on SHIMADZU LC-20AP instrument with original column. All final compounds were characterized by ^1^H NMR and HRMS. ^1^H NMR spectra were recorded on Bruker AVANCE III 500 MHZ (operating at 500 MHz for ^1^H NMR), chemical shifts were reported in ppm relative to the residual CDCl_3_ (*δ* 7.26 ppm ^1^H), DMSO-*d*_6_ (*δ* 2.50 ppm ^1^H) or Methanol-*d*_4_ (*δ* 3.31 ppm ^1^H), and coupling constants (*J*) are given in Hz. Multiplicities of signals are described as follows: s - singlet, d - doublet, t - triplet and m - multiple. High Resolution Mass Spectra were recorded on AB Triple 4600 spectrometer with acetonitrile and water as solvent.

#### 4.4.2 Cell lines and cell culture

The human cell line LNCaP was purchased from American Type Culture Collection. Cells were cultured according to the provider’s instructions and maintained at 37°C in a humidified atmosphere containing 5% CO_2_ in air. Cell lines were examined as mycoplasma free.

#### 4.4.3 Western blotting

1.5*×*10^5^ cells/ml were plated in 24-well plates and treated with compounds at the indicated concentrations for 24 h. Cells were collected and lysed in RIPA lysis buffer. Protein concentration was quantified by BCA. Antibodies used in this study include Androgen Receptor antibody (#5153S, Cell Signaling Technology), Anti-Rabbit IgG, HRP-linked (#7074S, Cell Signaling Technology) and GAPDH antibody (#8884 S, Cell Signaling Technology). Blots were imaged in Azure 300. ImageJ software was used to quantify Densitometry.

## 5 Data availability

The PROTAC-DB dataset are publicly available and can be accessed at http://cadd.zju.edu.cn/protacdb/. Both the PROTAC-ZL dataset and all preprocessed data used in this study for direct model training and evaluation are publicly available at https://github.com/LiZhang30/AiPROTAC.

## 6 Code availability

Our codes are available at https://github.com/LiZhang30/AiPROTAC.

## 7 Author contributions

L.Z. and X.L. collected and preprocessed the datasets. L.Z., M.W., X.L., X.C., and Y.L. designed the model and conducted all evaluation experiments. W.T., X.W., and K.Z. assisted in evaluation experiments. L.Z., X.L., and X.C. conducted case studies. R.S., C.R., and X.L. conducted biochemical experiments and analyzed all results. L.Z., M.W., X.C., X.L., R.S., and C.R. wrote and revised the manuscript. The study was supervised by X.Y., M.W., X.L., and X.C.

